# Expanded representation and function of PDF neurons in the mosquito circadian network

**DOI:** 10.64898/2026.11.19.767825

**Authors:** Linhan Dong, Anna V. Plank, Laura B. Duvall

## Abstract

Female *Aedes aegypti* mosquitoes can transmit pathogens when they bite humans, their preferred hosts, and this host-seeking drive is tightly gated by time of day. Daily rhythms depend on the molecular clock and communication among clock neurons, yet mosquito circadian network organization remains poorly understood. Using a mosquito-specific PERIOD antibody, we mapped the putative clock network of *Aedes aegypti* and found that neurons expressing the neuropeptide Pigment-Dispersing Factor (PDF) make up a much larger proportion of clock neurons compared to other insects, indicating a divergent network organization. PDF neurons are required for normal rhythms across insects, but whether their importance scales with their abundance in the network is unknown. We therefore tested the functional role of this expanded PDF neuron population in mosquitoes: we show that electrically silencing *Aedes aegypti* PDF neurons abolished the separation of daily activity into morning and evening peaks and led to misaligned clock protein rhythms across circadian neurons, even under entraining light-dark cycles. Thus, in a network built around an unusually large PDF population, PDF neuron activity plays an outsized role in behavioral timing and network synchrony.

---

Female mosquitoes require blood nutrients to develop eggs, and several species, including *Aedes aegypti*, specialize in finding and biting humans, making them particularly dangerous disease vectors^1,2^. Like many behaviors, mosquito host seeking and biting are gated by time of day, but this timing varies by species. Even among the small number of human-preferring species, rest-activity rhythms and biting times differ markedly. The African malaria mosquito, *Anopheles gambiae*, is mostly active at night, whereas the yellow fever mosquito, *Aedes aegypti*, is mostly active during the day^3^. Although light generally promotes biting in *Aedes aegypti*, biting is not uniform across the light phase of the day^4^. Instead, field and laboratory studies show a bimodal pattern, with host seeking peaking around dawn and dusk and dropping at midday^5–8^. This “temporal partitioning” of daytime activity may help mosquitoes avoid desiccation during hotter, drier hours of the day^9^, a strategy shared with the fruit fly *Drosophila melanogaster*, which shows a similar bimodal pattern with a midday “siesta”^10–12^.

The midday siesta, like other changes in activity within the light phase, depends on a functional circadian clock^10,13^. In *Drosophila*, individual loss of many major components of the transcription-translation feedback loop (TTFL) that make up the molecular circadian clock abolishes the siesta^13,14^. Similarly, in *Aedes aegypti*, mutants of the clock gene *cycle* show uniform activity throughout the day^15^. However, coherent behavioral rhythms require more than cell-autonomous clocks: they also depend on interactions among TTFL-expressing clock neurons. A key mediator of these interactions is the neuropeptide Pigment-Dispersing Factor (PDF), which regulates circadian rhythms in many insect species^16–20^. Extensive work in *Drosophila* has shown that PDF is critical for synchronizing molecular rhythms across clock neuron subgroups under constant conditions^21–23^.

Among the roughly 120 clock neurons per hemisphere in *Drosophila*^24^, only eight express PDF, divided evenly into two subsets: four small ventrolateral neurons (sLNvs) and four large ventrolateral neurons (lLNvs)^25^. Ablating or silencing these PDF+ cells causes defects in rhythmic behavior and molecular synchronization under constant conditions that resemble PDF null mutants. Under light-dark (LD) cycles, however, the bimodal activity pattern is largely preserved^16,26–28^. In many insect species, PDF neurons typically form a small population between the optic lobe and the central brain, with arborizations in the optic lobe and projections to the dorsal brain^29–31^. *Aedes aegypti* has PDF neurons in the same anatomical locations and with similar projection patterns, but in greater number^8,32^. This suggests a network organization that diverges from *Drosophila* and raises the possibility that PDF neurons play a broader functional role in mosquitoes.

Here, we generated a mosquito-specific antibody against PERIOD, a core component of the molecular clock, to profile putative clock neurons in *Aedes aegypti*, and used recently established genetic tools to electrically silence the PDF neurons and disrupt their communication. We find that PDF neurons make up a much larger fraction of the putative clock network in *Aedes aegypti* than in other insects studied, and that these neurons have an expanded functional role in behavioral regulation. Silencing PDF neurons abolishes the temporal partitioning of activity into morning and evening peaks, even under entraining LD cycles. These results suggest that, despite conserved PER dynamics, circadian networks can diverge across insect lineages in the abundance and weighting of core neuronal subpopulations, changing how much rhythmic behavior depends on individual contributions from these groups.

## Results

### PDF neurons make up about 40% of the putative mosquito circadian network

To survey putative clock neurons in *Aedes aegypti*, we generated a polyclonal antibody against the full-length *Aedes aegypti* PERIOD (PER) protein, a core component of the TTFL. Based on transcriptional evidence and earlier studies using *Drosophila* PER antibodies^32,33^, we examined clock neurons at zeitgeber time (ZT) 22 and consistently observed PER-immunoreactivity (PER-ir) in distinct neuron clusters (**Figure 1A–C**). Overall, PER-expressing neurons are found in two regions of the brain: the ventrolateral region (between the optic lobe and the central brain), and the dorsal region (**Figure 1A**). In the ventrolateral region (**Figure 1B**), the dorsal large ventrolateral neurons (lLNv-ds) sit directly anterior to the small ventrolateral neurons (sLNvs), and a separate group, the ventral lLNvs (lLNv-vs), sits lower. Co-staining with a PDF antibody shows that all lLNv-d and lLNv-v neurons are PDF+, whereas the sLNv group is mixed, with roughly one third of these neurons lacking PDF (**Figure 1D and E**). Because the sLNv group contains only PDF+ neurons in *Drosophila*, where the naming convention was established, we refer to the PDF+ neurons in this group simply as sLNvs and to the PDF-neurons as PDF-sLNvs. In the dorsal brain, a small group of neurons is found more medial and anterior, and a larger group is found more lateral and posterior (**Figure 1C**). We refer to these groups as the anterior dorsal neurons (DNas) and the posterior dorsal neurons (DNps), respectively. The dorsal projections of the sLNvs pass close to some DNps, suggesting potential communication (**Figure 1F**). **Figure 1G** summarizes the anatomical features of the putative clock neurons. In total, we report ∼43 putative clock neurons (42.8 ±0.7) across six subgroups per hemisphere (**Figure 1H**). PDF+ neurons account for ∼39% of this population, a strikingly high proportion compared with other insect circadian networks characterized with similar immunostaining-based methods, such as bees, ants, and aphids, where PDF+ neurons typically make up less than 10% (**Figure 1I**)^19,34–36^. This low proportion also appears to hold in *Drosophila*, where clock neurons have been labeled and counted using a wider range of methods than in other species, including genetic markers and connectomics^24,37,38^. Notably, the *Drosophila* DN3 group consists of small somas with dense anatomical overlap and its estimated size increased as immunostaining and genetic drivers were used more extensively^24^ but we did not observe a comparable group in *Aedes aegypti* with PER immunostaining. Overall, these anatomically distinct PER+ subgroups in the mosquito suggest distinctive functions and downstream targets, and the abundance of PDF+ neurons may indicate an expanded role for both the PDF peptide and PDF neurons in regulating behavioral timing.

**Figure 1.**
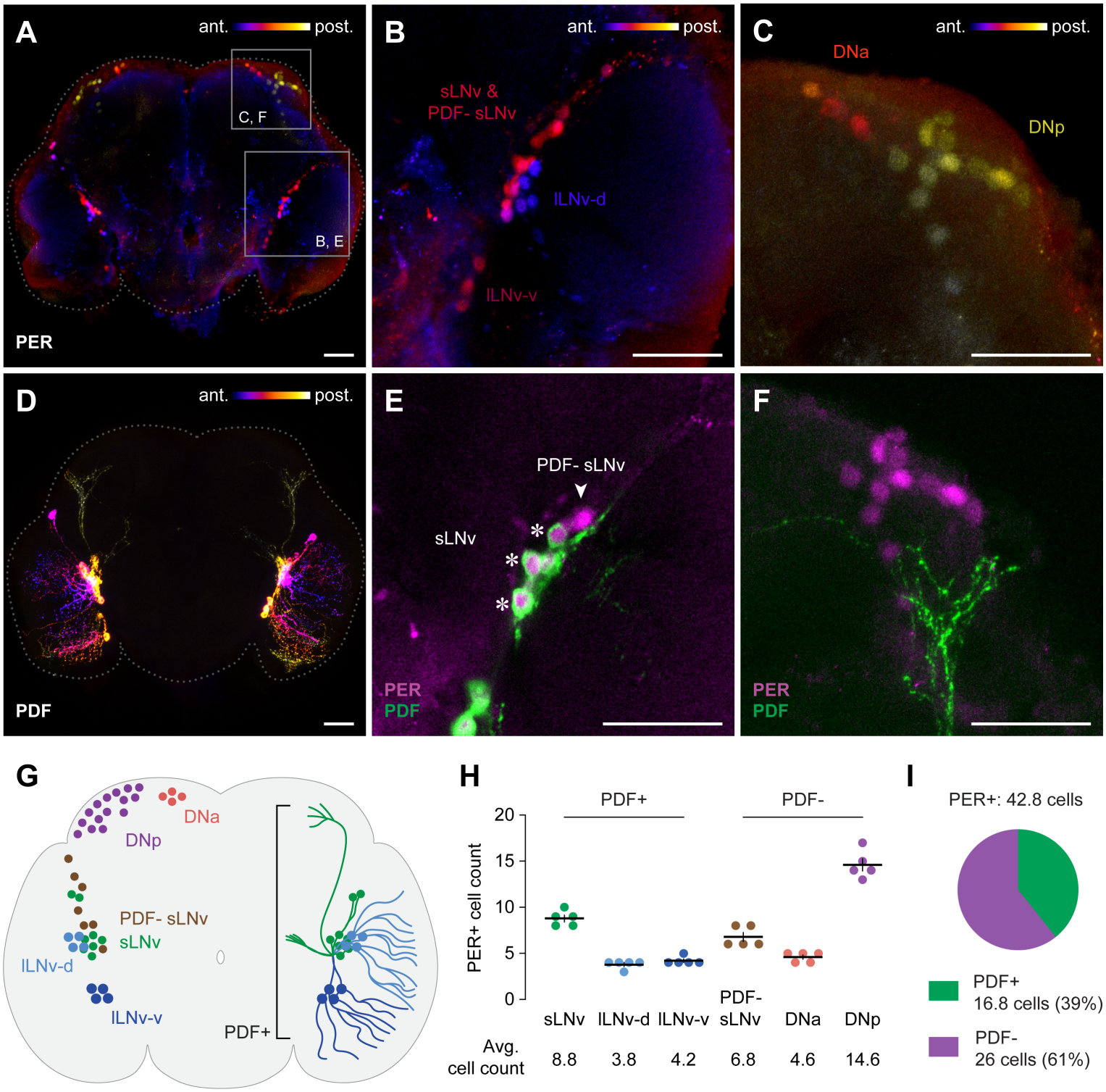
The putative mosquito circadian network contains a large proportion of PDF neurons. (A–C) Representative immunohistochemical staining of PER in female *Aedes aegypti* whole brain (A), ventrolateral region (B), and dorsal region (C). Maximum projection of relevant z-stacks. Scale bar: 50 μm. For (A–D), signal from a single channel is colorized according to anterior-posterior Z-slices. (D) Representative immunohistochemical staining of PDF in female *Aedes aegypti* whole brain. Maximum projection of relevant z-stacks. Scale bar: 50 μm. (E) Representative co-staining of PER and PDF in the ventrolateral region. Single optical section. Asterisks indicate PDF+ sLNvs and arrow indicates a PDF-sLNv. Scale bar: 50 μm. (F) Representative co-staining of PER and PDF in the posterior dorsal region. Maximum projection of relevant z-stacks. Scale bar: 50 μm. (G) Schematic of the *Aedes aegypti* putative circadian network. Left hemisphere shows the six PER+ neuron groups, and right hemisphere shows the PDF+ subsets and their projection pattern. (H) Cell count of PER+ neurons across subgroups (n = 5 brains). Cell counts are from one hemisphere. Bars and error bars show mean and SEM. (I) Average percentage of PDF+ neurons in the putative circadian network (n = 5 brains).

### PERIOD protein dynamics in the mosquito circadian network are conserved

Next, we surveyed the temporal dynamics of PER expression and localization in *Aedes aegypti* across four timepoints under a 12:12 h LD cycle. All putative clock cells in **Figure 1** showed daily PER oscillations (**Figure 2A–C**). PER-ir was detectable at ZT16, ZT22, and ZT4, but not at ZT10 (**Figure 2A–C**). To quantify PER oscillation in the PDF+ subgroups, we measured fluorescence intensity in the soma, which is marked by PDF immunoreactivity (PDF-ir) at all timepoints (**Figure 2A**). PER intensity peaks at ZT22 and is higher in the sLNvs than in the two lLNv groups (**Figure 2D**). Because the dorsal neurons and PDF-sLNvs lack clear secondary cellular or anatomical markers, we counted PER+ cells in these groups blind to condition. These counts show a similar oscillation, with no observable PER signal at ZT10 (**Figure 2E and F**). Overall, PER protein abundance closely matches the *Drosophila* pattern^39,40^, including the previously reported delay between *per* mRNA and protein (**Figure 3A**)^33,41^. Whole brain images also indicate PER oscillation in non-neuronal cells, similar to glial clock patterns reported in *Drosophila* (**Figure 2A**)^40,42,43^.

**Figure 2.**
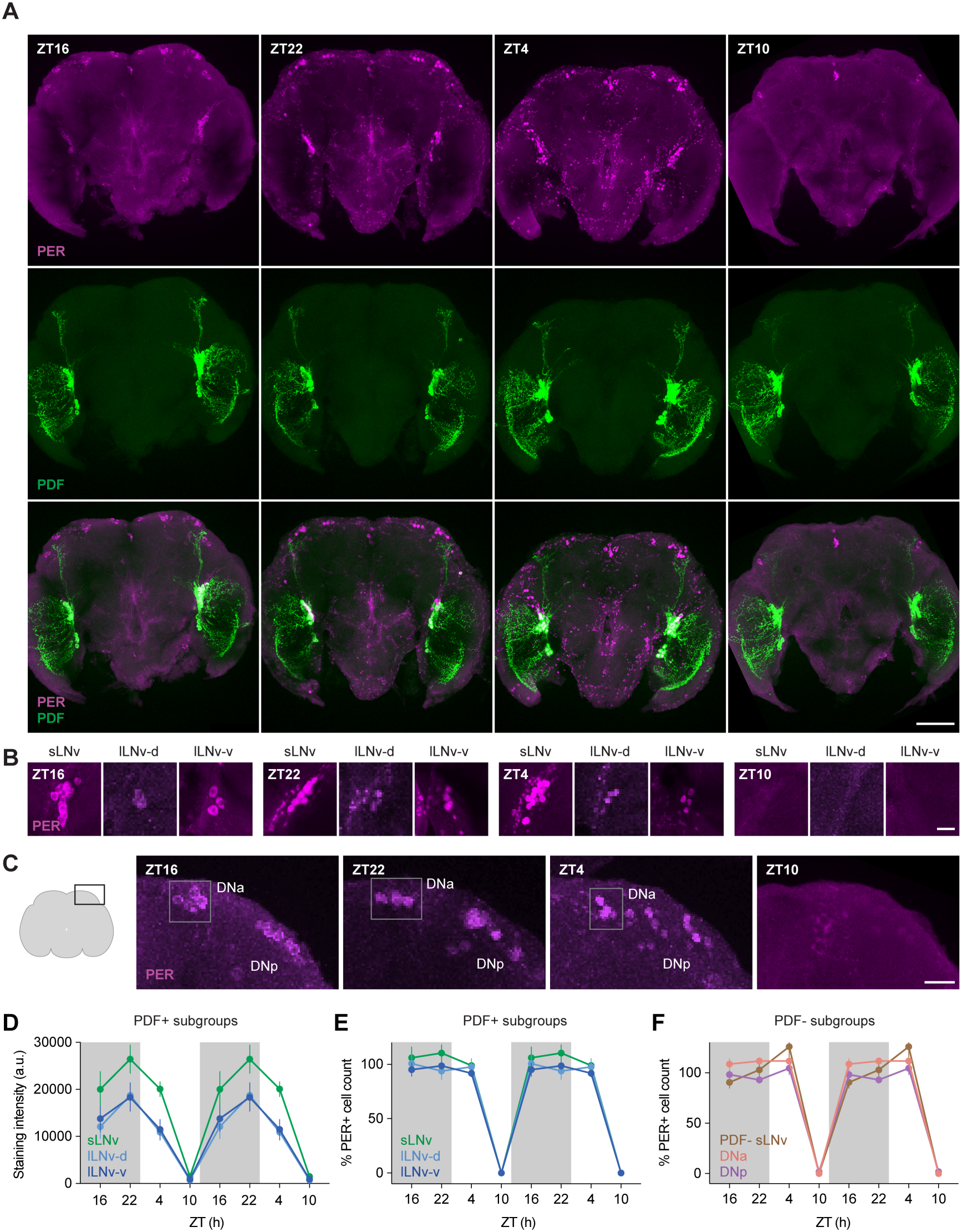
Putative circadian neurons exhibit daily oscillation in PERIOD expression. (A) Representative co-staining of PER and PDF across an LD cycle (ZT16, 22, 4, and 10). Maximum projection of relevant z-stacks. Scale bar: 100 μm. (B and C) Representative staining of PER in the ventrolateral region (B) and the dorsal region (C) across an LD cycle (ZT16, 22, 4, and 10). Maximum projection of relevant z-stacks. Scale bar: 20 μm. (D) PER staining intensity in the PDF+ neuronal subgroups (n = 6, 7, 7, and 7 brains at ZT16, 22, 4, and 10, respectively). Staining intensity reflects averaged measurements of somas within each subgroup (see Methods). (E and F) PER+ cell count in the PDF+ (E) and PDF-(F) neuronal subgroups (n = 6, 7, 7, and 7 brains at ZT16, 22, 4, and 10, respectively). Cell counts are from one hemisphere and normalized to cell count averages shown in **Figure 1H**. Data in (D–F) are double-plotted for visual continuance. Bars and error bars in (D–F) show mean and SEM. White and grey shading in (D–F) represent light and dark phases.

**Figure 3.**
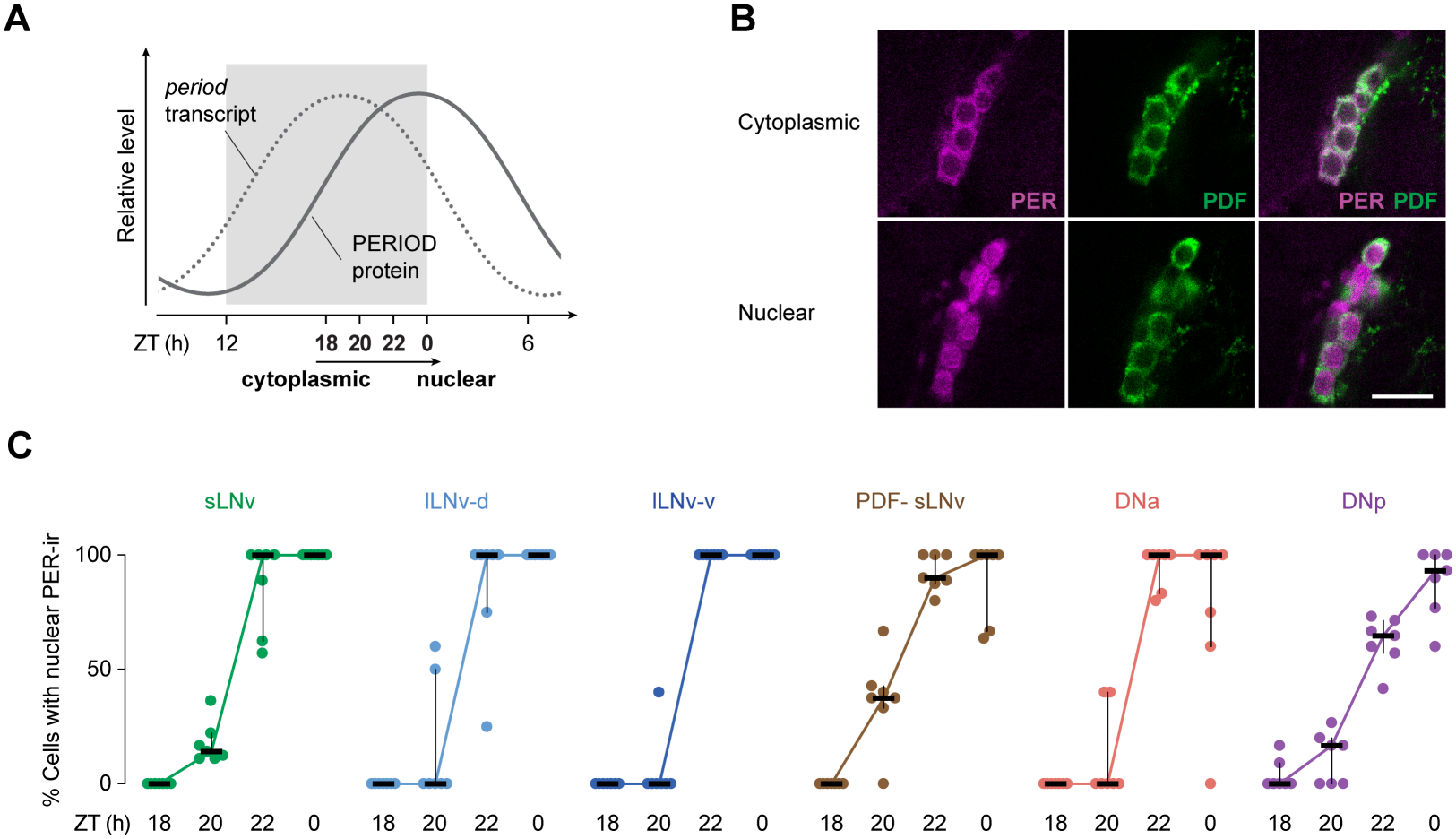
The clock protein PERIOD accumulates in the nucleus in a rhythmic manner in the mosquito. (A) Schematic of *per* mRNA and PER protein levels across the day. White and grey shading represent light and dark phases. (B) Representative PER and PDF staining in the sLNvs showing cytoplasmic PER-ir (ZT18) and nuclear PER-ir (ZT22). Maximum projection of relevant z-stacks. Scale bar: 20 μm. (C) Percentage of neurons with nuclear distribution of PER-ir across PER+ neuronal subgroups (n = 7 brains for all subgroups and timepoints). Bars and error bars show median and interquartile range. See also **Figure S1**.

Beyond protein abundance, PER also changes in subcellular distribution over time, a shift that is central to its function in the molecular clock. PER-ir is annular at ZT16 and solid at ZT22 (**Figure 2A–C**), typical of its translocation pattern in *Drosophila*, where PER first accumulates in the cytoplasm before entering the nucleus to inhibit the CLOCK-CYCLE transcriptional activator complex, shutting off *per* transcription and closing the negative feedback loop at the core of the molecular clock^44,45^. To investigate this translocation with finer temporal resolution, we compared PER-ir at additional timepoints (ZT18, 20, 22, and 0/24) to capture the nuclear entry of PER (**Figure 3A**). Across these timepoints, we identified two distinct patterns of subcellular distribution of PER-ir: cytoplasmic and nuclear (**Figure 3B**). In all six PER+ subgroups, PER enters the nucleus around ZT20 and PER-ir is nearly exclusively nuclear by ZT0/24 (**Figure 3C**, see also **Figure S1**). This suggests that nuclear accumulation of PER marks the repressive phase of the mosquito molecular clock, as in *Drosophila*, and that the timing of PER abundance and its subcellular localization are highly conserved.

### Silencing PDF neurons disrupts daily partitioning of activity under LD cycles

Coherent circadian rhythms require communication between clock neuron subgroups and synchronized molecular oscillations. In *Drosophila*, both the PDF peptide and PDF neurons are critical for behavioral rhythmicity and molecular syn-chrony among clock neuron subgroups under free-running conditions^16,22,23,46^. Like *pdf*^*01*^-null mutants, flies with electrically silenced PDF neurons lose circadian rhythms and show phase misalignment among PDF-neuronal subgroups^26–28,47^. To ask whether the expanded representation of PDF neurons in the mosquito clock network is accompanied by greater functional importance, we used a genetic approach to electrically silence them. We previously established a *pdf* mutant carrying a QF2 insertion^8^. Using the Q-binary expression system, we showed that, when used as a heterozygous driver, this line labels PDF cells in the brain with high specificity^8^. Here, we used the *pdf-QF2* line in combination with a recently established QUAS effector line to drive expression of the inward-rectifying K^+^ channel, Kir2.1, in PDF neurons^48^, and we confirmed expression using eGFP-tagged Kir2.1 (**Figure S2**).

To test whether Kir2.1 expression in PDF neurons causes behavioral defects, we measured spontaneous locomotor activity rhythms under 12:12 h LD cycles. Both parental lines showed normal bimodal activity patterns with a midday siesta around ZT7 (**Figure 4A and B**, see also **Figure S3A**). In contrast, *pdf > Kir2*.*1* animals lost this bimodal pattern and their midday siesta was largely abolished. This suggests that normal electrical activity of PDF neurons is required to suppress midday activity even under entraining LD cycles. This constant daytime activity pattern resembles the phenotypes of clock mutants in *Drosophila*, and *cycle* knock-outs in *Aedes aegypti* (**Figure S3A**)^13,15^. The lack of activity variation within the photophase or scotophase can be measured by the slope of anticipatory activity^49^. Because *Aedes aegypti* do not show anticipatory activity before lights-on, we measured the anticipation slope for the evening peak, which decreased when PDF neurons were silenced (**Figure 4C**). Although darkness suppresses activity in *Aedes aegypti*, evening activity usually extends for about 30 minutes after lights-off^7,8,50^. The *pdf-QF2* knock-in is itself a *pdf* mutant, and this extension was shortened in the heterozygous *pdf-QF2* parental controls compared to QUAS controls. However, it was almost completely abolished in *pdf > Kir2*.*1* animals, a stronger effect than in either parental line (**Figure 4D**). Flattening of evening anticipation and lack of activity extension after lights-off seen in *pdf > Kir2*.*1* animals are also observed in *cycle* knock-outs (**Figure S3B and C**). Overall, silencing PDF+ clock neurons phenocopies loss of clock gene function for these measures, indicating that, without normal PDF neuron activity, mosquitoes cannot partition their activity into a bimodal pattern.

**Figure 4.**
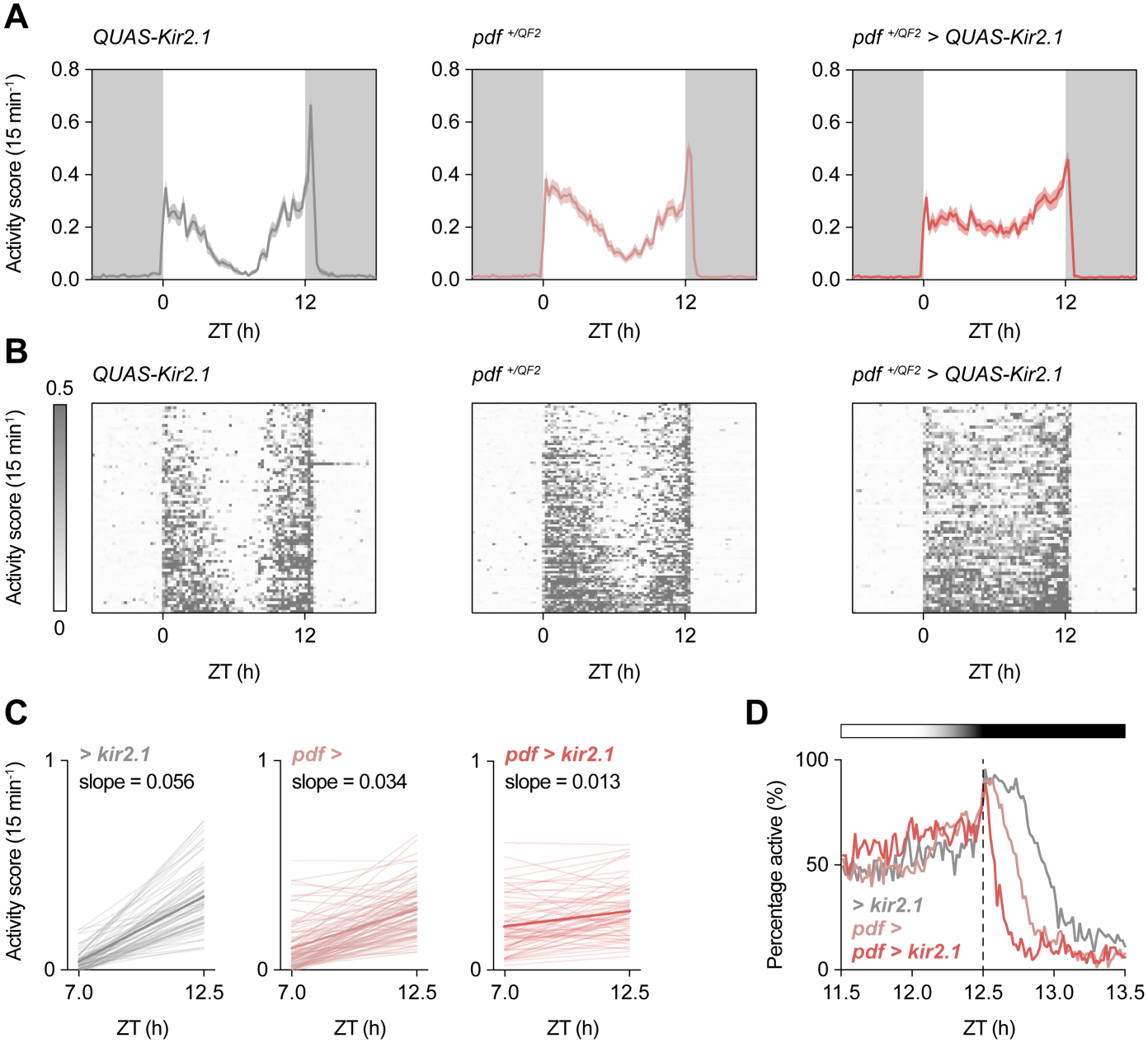
Electrical silencing of PDF neurons disrupts temporal partitioning of *Aedes aegypti* activity. (A) Locomotor activity rhythms of female mosquitoes under LD cycles. For all panels, data shown is for the third day under LD cycles. For all panels, n = 71, 91, and 66 animals for *> Kir2*.*1, pdf >*, and *pdf > Kir2*.*1*, respectively. White and grey shading represent light and dark phases. Line and shading show mean and SEM. (B) Individual actograms under 12:12 h LD cycles. Each row represents a single individual. (C) Linear regressions of activity score before lights-off. Thin lines show individuals and thick lines show group average. (D) Percentage of active animals around lights-off. Light and dark shading represents relative light intensity and dotted line represents lights-off. See also **Figure S2–4**.

To test whether the increased midday activity also reflects changes in rest, we analyzed rest using a 10-min inactivity threshold^51^. During the day, *pdf > Kir2*.*1* mosquitoes showed significantly less total rest, fewer rest bouts, and shorter average rest duration (**Figure S4A–C**). At night, they showed more total rest and longer average rest duration (**Figure S4D–F**). Because rest bouts were reduced across both day and night, we reasoned that electrically silencing PDF neurons may impair rest initiation. These results indicate that the loss of bimodal activity partitioning reflects both mistimed activity and disrupted daytime rest.

### Silencing PDF neurons disrupts clock synchronization in DNp clock neurons under LD

In *Drosophila*, the role of PDF and PDF neurons in synchronizing the circadian network is well established under constant darkness (DD). Silencing PDF neurons or disrupting the *pdf* gene misaligns molecular oscillations among clock neuronal subgroups, which provides a mechanistic explanation for the loss of circadian rhythmicity under DD^22,26–28,47^. Because silencing PDF neurons also disrupted behavioral rhythms under LD in *Aedes aegypti*, we asked whether it misaligns clock oscillations under entraining conditions by measuring PER expression at its peak and trough (ZT22 and ZT10). PER signal was generally higher at ZT22 than at ZT10 across the ventrolateral, DNa, and DNp regions, (**Figure 5A–C**), but exact oscillation patterns showed subgroup-specific differences. Among PDF+ neurons, PER intensity differed between ZT22 and ZT10 in the sLNvs of both *pdf > Kir2*.*1* animals and parental controls (**Figure 5D**). In the two lLNv groups, PER oscillation was dampened to varying degrees in both *pdf >* heterozygous controls and *pdf > Kir2*.*1* mosquitoes (**Figure S5**), suggesting that this defect reflects the loss of one copy of *pdf* in the *pdf-QF2* knock-ins^8^. Among PDF-subgroups, blinded scoring of PER-ir showed signal at ZT22 but not at ZT10 in the PDF-sLNvs and the DNas (**Figure 5E and F**). However, in the DNps, which lie near the dorsal terminals of PDF neurons (**Figure 1F**), about five neurons in *pdf > Kir2*.*1* mosquitoes showed signal at ZT10, when PER is normally undetectable (**Figure 5G**). This suggests that synchronization of these DNps requires electrical activity of PDF neurons even under LD conditions. Since only about a third of all DNps were out of phase, the DNp region may contain functional and anatomical subdivisions that we cannot yet resolve. Overall, silencing PDF neurons affects clock synchronization under LD in a subgroup-specific manner, most clearly in the DNps.

**Figure 5.**
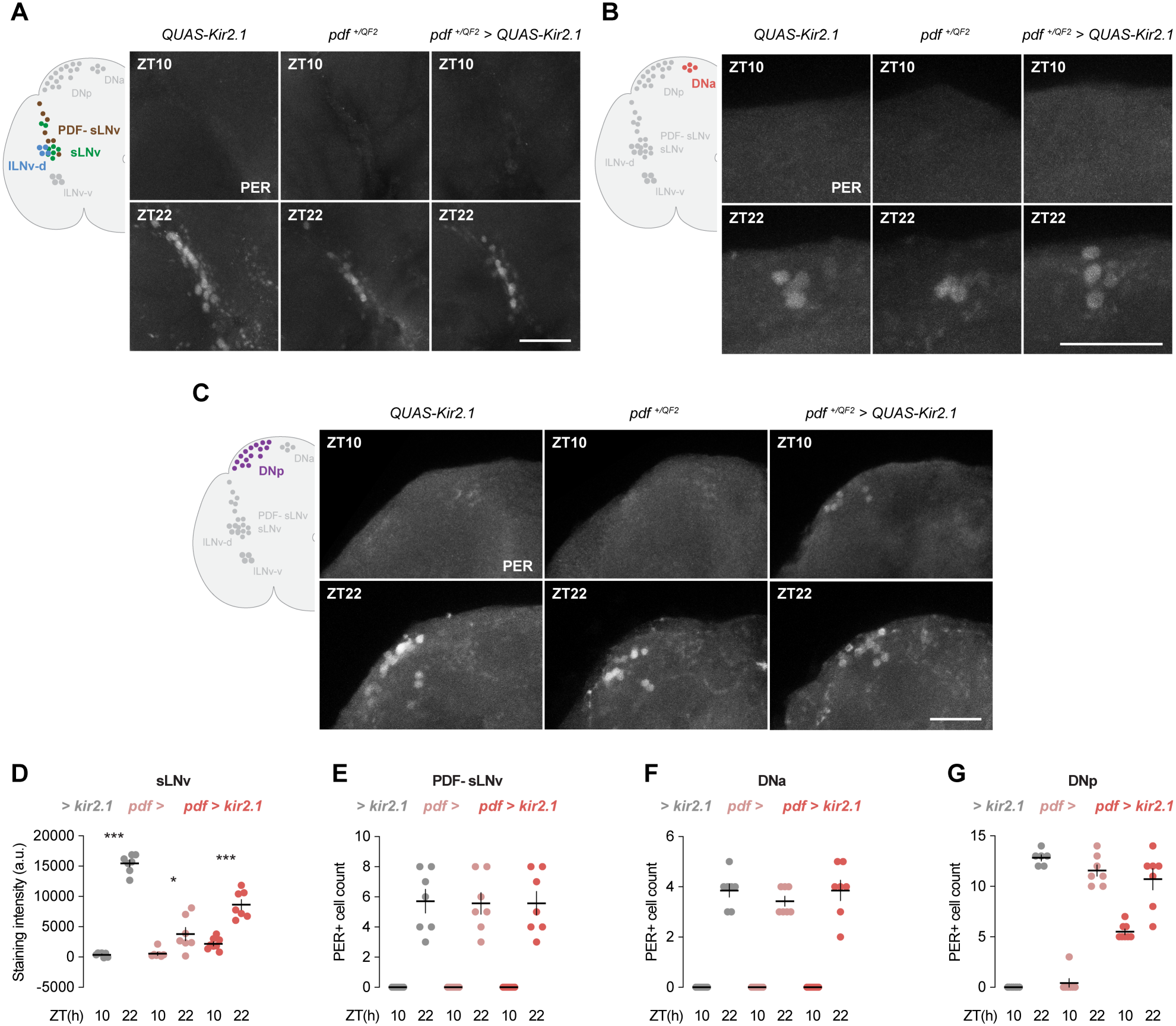
Electrical silencing of PDF neurons disrupts PERIOD oscillation in DNp neurons. (A–C) Representative immunohistochemical staining of PER in the ventrolateral regions (A), the DNa regions (B), and the DNp regions (C). For all panels, n = 6–8 brains for all genotypes and timepoints. Maximum projection of relevant z-stacks. Scale bar: 50 μm. (D) PER staining intensity in the sLNvs. See also **Figure S5**. Staining intensity reflects averaged measurements of somas within each subgroup (see Methods). Mann-Whitney tests. (E–G) PER+ cell counts in the PDF-sLNvs (E), DNas (F), and DNps (G). Cell counts are from one hemisphere per brain. Line and error bars in (D–G) show mean and SEM and dots show individual animals. *P < 0.05; ***P < 0.001. Exact n and P values are provided in **Table S1**. See also **Figure S5**.

## Discussion

### Neuronal control of mosquito behavioral timing

Disease vector mosquitoes like *Aedes aegypti* are remarkably precise at finding and biting humans^52^. Host attraction relies on multimodal sensory pathways that encode robust host-seeking strategies, driven by the mosquito’s strong reproductive need for blood^53,54^. However, a less appreciated aspect of biting behavior is that host-seeking drive is strongly modulated by time of day and the circadian clock. In both diurnal and nocturnal species, biting is not uniform across the light or dark phase, but is instead concentrated in specific windows usually occurring around dawn and/or dusk^9,55^. Partitioning activity into these distinct peaks requires the endogenous circadian clock, and knocking out a core TTFL component abolishes the bimodal activity rhythm in *Aedes aegypti*^15,56^.

However, clock gene mutants disrupt the molecular clock in every cell, so, while they demonstrate that the clock is required, these manipulations cannot reveal which neurons drive behavioral timing. Answering these questions requires manipulating the activity of defined neuronal populations. This is made possible by the recent development of genetically encoded tools in mosquitoes, including the electrical silencing effector line *QUAS-Kir2*.*1*, recently applied in peripheral gustatory neurons^51^. Our work extends the application of these tools to the central brain and the circadian system. By combining our *pdf*-QF2 driver line with Kir2.1, we silenced PDF neurons without directly manipulating the molecular clock^8^. PER continues to oscillate in the sLNvs under silencing (**Figure 5D**), indicating that the molecular clock remains functional. Kir2.1-mediated silencing is expected to suppress the release of action-potential dependent neuronal signals from these cells, and because electrical silencing primarily affects neuronal activity, comparing these phenotypes with those of *pdf* mutants can distinguish peptide-specific from neuron-wide contributions.

Silencing PDF neurons reproduced key features of clock gene loss including loss of siesta, flattened evening anticipation, and early termination of evening activity, and it desynchronized a subset of DNp clock neurons. These results indicate that PDF neuron activity is needed to keep the clock network synchronized and to translate that timing into a bimodal activity pattern even under LD cycles. These defects indicate that, without PDF neuron activity, behavior is driven more by light-dark transitions than by internal timing, compromising coherent activity-rest regulation, which may lead to timing shifts in host seeking and biting. Although the downstream targets that mediate these behavioral effects, such as the DNps near the PDF terminals, remain to be characterized, our findings establish an important role for PDF neurons in coordinating circadian network function and behavioral timing. The development of new driver and effector lines will make it possible to directly test which neuronal populations control the timing of specific behaviors, including host-seeking and biting.

### Divergent anatomical and functional organization of the mosquito circadian network

Animal circadian clocks are complex neuronal networks that coordinate molecular oscillations and drive many physiological and behavioral processes. The anatomical and functional diversity of clock neurons has been dissected most thoroughly in *Drosophila melanogaster*. Early work on clock neuron sub-groups suggested direct behavioral output from morning and evening oscillators and a principal pacemaker^16,46,57,58^, but recent studies have identified more complex networks and alternative drivers of rhythmic behavior^59–61^. This heterogeneity extends to how the network is coordinated; pacemaker subgroups rely on phase synchronized clock oscillations to generate nonsynchronous calcium rhythms and distinct rhythmic outputs^62,63^. Synchronization depends on both PDF and PDF neurons, and without normal PDF signaling, behavioral, molecular, and neuronal activity rhythms are severely disrupted under constant conditions^16,21,22,26,27,57,62^. However, under LD conditions, PDF disruption only moderately affects bimodal activity patterns and molecular oscillations in *Drosophila*^26,27,47^. This suggests that the *Drosophila* clock can use alternate pathways to synchronize molecular rhythms when external zeitgebers are present.

Although mosquitoes and fruit flies are both true flies (Diptera), their genetic clocks differ in terms of molecular composition of the clock feedback loop, transcript dynamics, and clock gene function^33,55,64,65^. Here, we show a mosquito circadian network that also diverges from *Drosophila* both anatomically and functionally. Anatomically, these mosquitoes have more than twice as many PDF neurons (17 per hemisphere in *Aedes aegypti*, 8 per hemisphere in *D. melanogaster*), and PDF neurons make up ∼40% of PER+ neurons that we identified in *Aedes aegypti*^24^. It is important to note that the characterization of *Drosophila* clock neurons accumulated extensive research using immunostaining as well as genetic labeling. However, characterization of other insect circadian networks using similar approaches methods also shows that these mosquitoes have a larger number and proportion of PDF neurons compared to honeybees, pea aphids, and ants^19,34–36^. Functionally, silencing PDF neurons abolishes the siesta and disrupts evening activity even under LD conditions in *Aedes aegypti*, whereas in *Drosophila* LD cycles are sufficient to maintain the bimodal pattern even with disrupted PDF. The *pdf > Kir2*.*1* mosquitoes show a clock-mutant like phenotype, which differs from the PDF knock-out, where the morning peak was diminished but the evening peak was unaffected^8^. This implies that the activity of PDF cells and the signaling function of the PDF peptide do not produce identical behavioral effects. Collectively, these results show that the PDF-biased *Aedes aegypti* circadian network is both anatomically distinctive and functionally divergent from that of *Drosophila*, demonstrating that differential behavioral regulation can be evolved via differential clock neuronal organizations while preserving core molecular clock dynamics.

### Evolutionary implications of a PDF-biased clock network

The divergent organization of the mosquito clock network raises questions about the evolutionary significance of different neuronal architectures for circadian timekeeping. The prominence of PDF-expressing neurons in the *Aedes aegypti* clock may reflect a distinct strategy for coordinating daily activity patterns. PDF neurons are broadly implicated in light input and entrainment, so a PDF-biased network might increase the influence of environmental light cues on the timing of behavioral rhythms^59^. Consistent with this, light strongly affects activity in diurnal *Aedes aegypti*^4,8^ and recent work also suggests that the mosquito clock can be rapidly reset by environmental light, with a single alteration in the light cycle sufficient to induce a phase shift^66^. Although *Aedes aegypti* inhabits tropical and subtropical regions with relatively limited seasonal variation in photoperiod, its close association with humans exposes it to environments in which artificial lighting can alter natural light-dark cycles. This raises the possibility that strong light responsiveness may influence how mosquitoes time their activity in human-modified environments, where host availability may not follow natural daily light cycles^4,67^. Our recent work showed that female mosquitoes switch their temporal niche during a reproductive cycle^56^, further demonstrating that the timing of mosquito activity is influenced by both environmental and physiological factors. Comparative work across species with different degrees of human association could test whether coexistence with humans shaped the evolution of the *Aedes aegypti* clock and how it orchestrates activity patterns.

### Limitations of the study

The *pdf-QF2* genetic driver was previously generated as a knock-in/knock-out mutant, and heterozygotes show behavioral and anatomical effects^8^. These behavioral defects were more evident under DD conditions than LD conditions, suggesting that both copies of the *pdf* gene are required to maintain rhythmicity without external zeitgebers. To avoid confounding effects of the heterozygous driver, we therefore examined the effect of PDF neuronal silencing only under LD conditions where the heterozygotes show normal bimodal activity. Future drivers that do not disrupt *pdf* could allow additional studies under constant conditions. A second limitation is that our putative circadian network is based on antibody staining of the PER protein itself. As in *Drosophila*, the use of multiple genetic drivers and antibody labeling might reveal more clock neurons that are currently undetected^24^. Expanding the circadian genetic and molecular toolbox with additional clock gene drivers will be critical to advance our understanding of the mosquito TTFL and circadian network.

## Acknowledgements

We thank Olivia Martinez and Thomas Gabel for assistance with animal husbandry, Inneloy Polanco for assistance with locomotor activity recordings. We thank Trevor Sorrells and Anjali Pandey for generating and providing the *QUAS-Kir2*.*1* line. We thank Nils Reinhard and members of the Duvall lab for useful discussions and feedback on the manuscript. This work was supported by the following grants: NIGMS (R35 GM137888) (LBD).

## Author contributions

LD carried out or supervised all experiments and data analysis. AVP performed animal circadian entrainment, immunohistochemistry, confocal brain imaging, and image analysis with LD. LD, AVP, and LBD conceived the study, designed the figures, and wrote the paper.

## Supplemental information

**Table S1**. All raw data and statistical test details used for main and supplemental figures.

## Methods

### Resource availability

#### Lead contact

Requests for resources and reagents should be addressed to lead contact, Laura B. Duvall.

#### Materials availability

Polyclonal rabbit PERIOD antibody is available upon request.

#### Data and code availability

- All data used in main and supplemental figures are available in the supplementary materials.
- No original code was used in this study.
- Any additional information required to re-analyze the data reported in the paper is available from the lead contact upon request

#### Experimental model and study participant details

All mosquitoes used in this study are wild-type *Aedes aegypti* (Liverpool strain) or transgenic of the same background. The *pdf-QF2* line and the *QUAS-eGFP-Kir2*.*1* line were previously generated and crossed as heterozygotes to generate the *pdf > Kir2*.*1* cross^8,48^. Both parental lines were maintained as heterozygotes, and offspring of the two parental lines were screened during the larval or pupal stage for a dsRed (*pdf-QF2)* or eCFP (*QUAS-eGFP-Kir2*.*1*) eye marker, or both (*pdf > Kir2*.*1*), using a Leica fluorescence microscope. Animals were reared at 28°C with 70–80% humidity using a 12-h light and 12-h dark cycle. Eggs were hatched with 1 L of a hatch broth solution made with ground TetraMin Tropical Tablets or Tropical Flakes fish food in deoxygenated water. Larvae were fed using this same ground fish food. Upon eclosion, males and females were cohoused for at least 3 days and given *ad libitum* access to a 10% sucrose solution. For all experiments, female mosquitoes 4–14 days old were used. For passaging, co-housed females were blood-fed using defibrinated sheep blood (Quad5) warmed to 37°C with the Hemotek membrane feeding system. Eggs were collected on filter papers (Whatman), and egg papers were air-dried and stored at room temperature until used for hatching.

### Method details

#### Generation of the polyclonal PER antibody

Polyclonal rabbit anti-PER antibodies were generated by GenScript. Briefly, the *period* transcript variant sequence X4 (XM_021854837) was used as antigen sequence. The sequence is modified with a His-tag and this 1100 aa fragment was cloned into a pET30a vector and expressed in *E. coli* BL21 Star (DE3) competent cells. Exact antigen sequence is included in **Table S1**. Immunization was performed on two New Zealand rabbits, and purified antibody was verified against the antigen using ELISA.

#### Whole-mount immunohistochemistry

Prior to tissue fixation dissection, 4–11 day old co-housed females were cold-anesthetized and transferred into individual wells in a 12-well plate for circadian entrainment, with each well containing 1.5 mL of a 10% sucrose solution in 1% agarose supplemented with 1% of an antifungal agent (10% p-hydroxy-bensoic acid methyl ester in 95% Ethanol). These 12-well plates were then placed in a custom-built locomotor activity monitor “MozzieBox” for 3–4 days at 28°C. The setup and operation of the MozzieBox were previously described^56^. Briefly, the MozzieBox uses a RaspberryPi computer to set a light entrainment schedule inside of a light-proofed incubator (Heratherm, Thermo Fisher), and white LED lights inside the box provide lighting. All entrainment was done with a LD cycle of 12h: 12h, with a 30-min light transition period. For all staining procedures used in quantitative analysis (**Figure 2** and **Figure 5**), samples from all ZT timepoints were processed simultaneously from different ZT0 settings in the MozzieBox entrainment schedules. All animals were 7–14 days old at the time of fixation and dissection.

At the desired ZT time point, animals were transferred to 4°C.

For timepoints during the dark phase (ZT16, ZT18, ZT20, and ZT22), animals were transported in a light-tight container (BK5, Thorlabs), and were handled within 20 mins under a dim red light. A group of 9–12 mosquitoes were fixed in 4% paraformaldehyde in 1x PBS containing 0.5% Triton X-100 (PBS-T) for 2 hours and 15 minutes at 28°C, rocking on a nutator. Tubes containing animals collected in the dark phase were wrapped in tinfoil during this fixation step. After fixation, animals were washed 5 times with PBS-T (1x PBS, 0.5% Triton X-100) at room temperature. Brains were then dissected in the PBS-T, and then blocked in PBS-T supplemented with 5% Normal Goat Serum (NGS) overnight at 4°C. The brains were washed in PBS-T for 10 mins at room temperature and then incubated in a primary antibody solution (5% NGS in PBS-T supplemented with 1:500 mouse anti-PDF (DSHB, PDF C7) and/or 1:1000 rabbit anti-PER (this study) for all PER-staining except the brains in **Figure 1G–I**, which used 1:500 mouse anti-PDF and 1:500 rabbit anti-PER) for 3 days at 4°C. For the brains in **Figure S2**, the primary antibody solutions were made with 1:250 mouse anti-PDF (DSHB, PDF C7) and 1:500 chicken anti-GFP (Aves Labs, GFP-1020). Following primary antibody incubation, brains were washed 5 x 10 minutes in PBS-T at room temperature. Samples were then incubated overnight at 4°C protected from light in secondary antibodies solution (5% NGS in PBS-T supplemented with 1:500 of secondary antibody). The following secondary antibodies were used: goat anti-mouse-Alexa488 (Thermo Fisher Scientific, A-11001), goat anti-rabbit-Alexa647 (Thermo Fisher Scientific, A-21245), goat anti-rabbit-Alexa488 (Thermo Fisher Scientific, A-11008), goat anti-mouse-Alexa647 (Thermo Fisher Scientific, A-21236), and goat anti-chicken-Alexa488 (Thermo Fisher Scientific, A-11039). Prior to mounting, the brains were washed 3 x 10 minutes in PBS-T, remaining protected from light. Brains were then cleared and mounted in RapiClear (SunJin Lab) on glass slides using an imaging spacer (iSpacer 0.2mm, SunJin Lab). Mounted samples were sealed with super glue on the corners and nail polish around the edges, and incubated at room temperature for at least 30 minutes to achieve optical clearing prior to confocal imaging.

#### Confocal imaging

Mounted slides were imaged on a Zeiss LSM 880 laser scanning confocal microscope using a Plan-Apochromat 20×/0.8 M27 air objective controlled by Zeiss ZEN (Black edition) software. Images were acquired at 16-bit depth with dimensions of 2048 × 2048 pixels, a line average of 2, a scan speed of 8, and a z-step size of 2 μm. The pinhole was set to 1.0 AU for each channel. Representative images in **Figure 3B** were obtained with a C-Apochromat 40×/1.2 Korr FCS M27 objective, with a scan speed of 9. All images used for quantitative analyses were obtained in the same imaging session.

#### Locomotor activity assay

To assess locomotor activity in a controlled environment, co-housed females were loaded into the MozzieBox system in the same way as described above (see ***Whole-mount immunohistochemistry***). Plates were placed inside the MozzieBox at 28°C under continuous infrared illumination (940 nm LED strips), and the animals were subject to a LD cycle of 12h: 12h for 3 days, with a 30-min light transition period. Plate positioning within the box was randomized. During this time period, Raspberry Pi 5MP 1080p NoIR cameras captured images at 1-minute intervals. At the end of the experiment, images were compiled into a video using custom Matlab code and animals were tracked with the EthoVision (version 17) software to report the activity of individual mosquitoes. All code used for the operation of the MozzieBox locomotor activity assay and downstream data processing was previously described^56^.

#### Quantification and statistical analysis Image processing and analysis

All confocal images were processed and analyzed using Fiji (version 2.16.0/ImageJ 1.54p). All representative images showing quantitative signal intensities were displayed with no adjustments or uniform adjustments on brightness and contrast to enhance visibility using Photoshop (version 24.3.0). For all analyses, the right side of the brain was analyzed unless significant damage was observed, in which case the left side was used. All image analyses described below were blinded to the experimenter.

For cell count analyses (**Figures 1H–I, 2E–F, and 5E–G**), cells were manually counted in one hemisphere throughout the whole brain z-stack. In **Figure 2E–F**, values graphed are obtained by dividing the number of cells seen by the average count for each cell group shown in **Figure 1H**. For **Figure 1H and I**, brains collected at ZT22 were used.

For subcellular localization analysis in **Figure 3C**, the total number of cells was counted for each group as described above. Then, a second count was performed, tallying the number of cells appearing to have nuclear localization. Nuclear localization was defined as cells that lack an annular shape with lower nuclear intensity (see **Figure 3B**).

For staining intensity quantitative analysis, used in **Figure 2D, Figure 5D**, and **Figure S5**, PDF staining was used as a marker for cell soma boundaries. Polygon ROIs were manually drawn and used to measure PER staining intensity in the same z-slice. Three rectangular background measurements of a fixed size were taken from the same z-slice. Intensity was then calculated by subtracting the average background mean from the signal mean. In most cases, three cells from each cell group were analyzed and averaged (in some rare cases, only two lLNv-ds were observed, measured and averaged).

#### Locomotor activity analysis

All data processing for the MozzieBox locomotor activity assay was previously described^56^. Briefly, raw movement data in 1-min intervals were converted to an activity index on a 0-100 linear scale for evening termination analysis and rest analysis, or summed as 15-min bins for all other plotting or analysis. Evening activity anticipation analysis fits a simple linear equation through the 15-min binned activity data across ZT7–ZT12.5 in the third LD cycle, which was performed with GraphPad Prism (version 10.3.0). For evening activity termination analysis and rest analysis, inactivity is defined as activity index = 0. All rest analyses were performed with using the phase package (1.2.9) in R (version 4.2.3), with the rest definition as 10 minutes of inactivity. All data used in **Figure S3** were collected in a previous publication^56^, and raw data were reanalyzed and replotted here.

#### Statistical analysis

All statistical analyses were performed with GraphPad Prism (version 10.3.0). Nonparametric tests were used for all statistical comparisons, and methods and sample sizes are reported in the figure legends. All raw data used in main and supplemental figures, with accompanying test details and exact sample sizes, are included in **Table S1**.

## Supplemental figures

**Figure S1.**
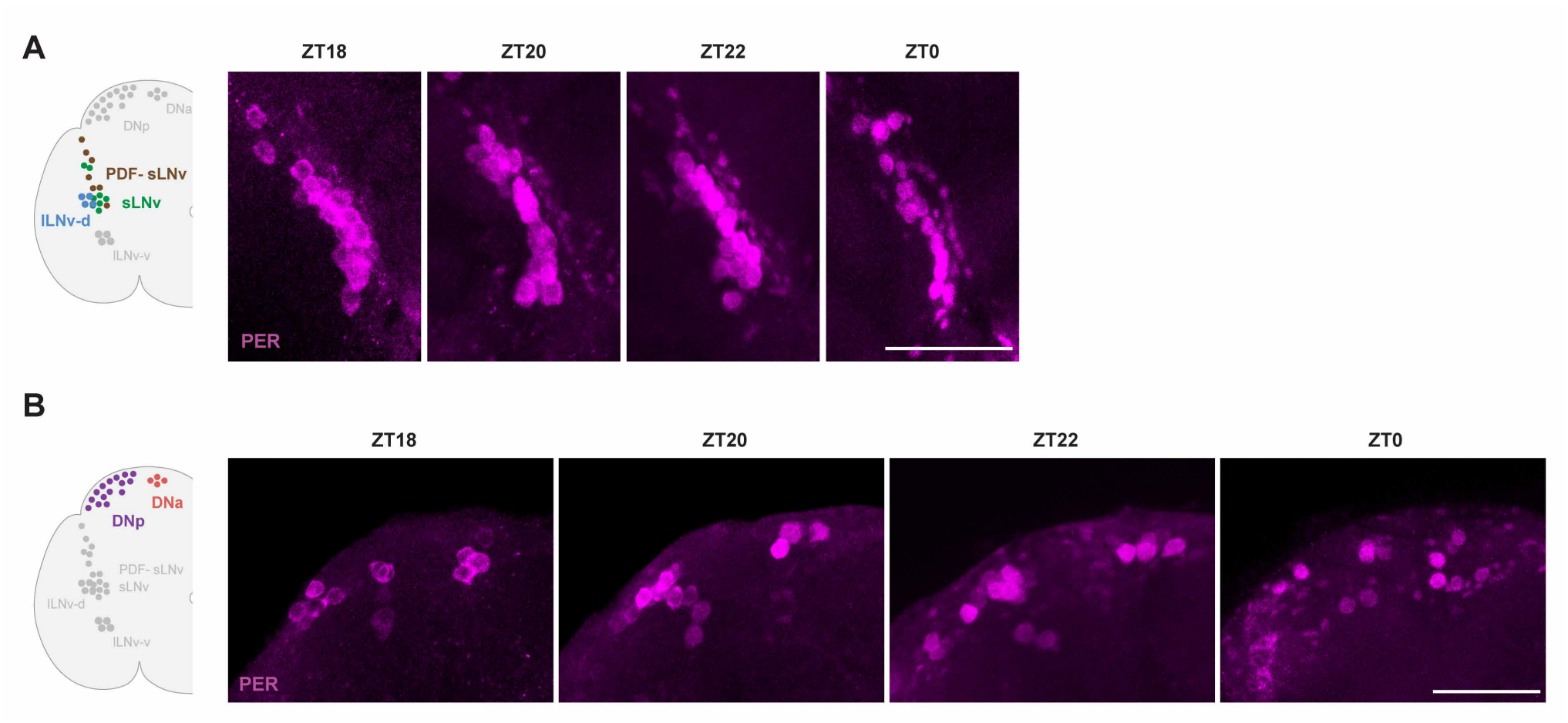
Subcellular distribution of PER during the scotophase in the ventrolateral region (A) and the dorsal region (B), related to Figure 3. Representative images. Maximum projection of relevant z-stacks. Brightness and contrast were adjusted across the four ZT timepoints to normalize for signal differences for better visualization of the subcellular localization. Scale bar: 50 μm.

**Figure S2.**
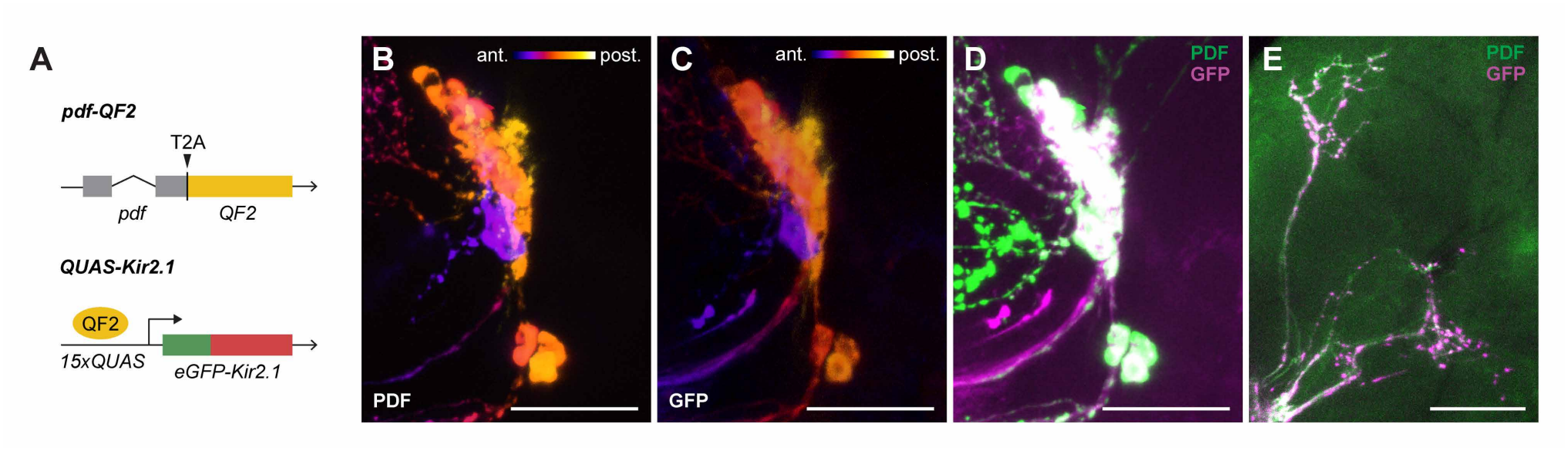
Characterization of the *pdf > Kir2*.*1* genetic construct in the mosquito brain, related to Figure 4. (A)Schematic of the *pdf-QF2* driver and *QUAS-Kir2*.*1* transgenic lines. (B)Representative PDF immunostaining of PDF cells in *pdf > Kir2*.*1* animals. For (B) and (C), signal from a single channel is colorized according to anterior-posterior Z-slices. (C)Representative GFP immunostaining of PDF cells in *pdf > Kir2*.*1* animals. (D)Representative co-staining of PDF and GFP of PDF cells in *pdf > Kir2*.*1* animals. (E)Representative co-staining of PDF and GFP in the PDF cell dorsal projections of *pdf > Kir2*.*1* animals. Images in (B–E) are maximum projection of relevant z-stacks. Scale bar: 50 μm.

**Figure S3.**
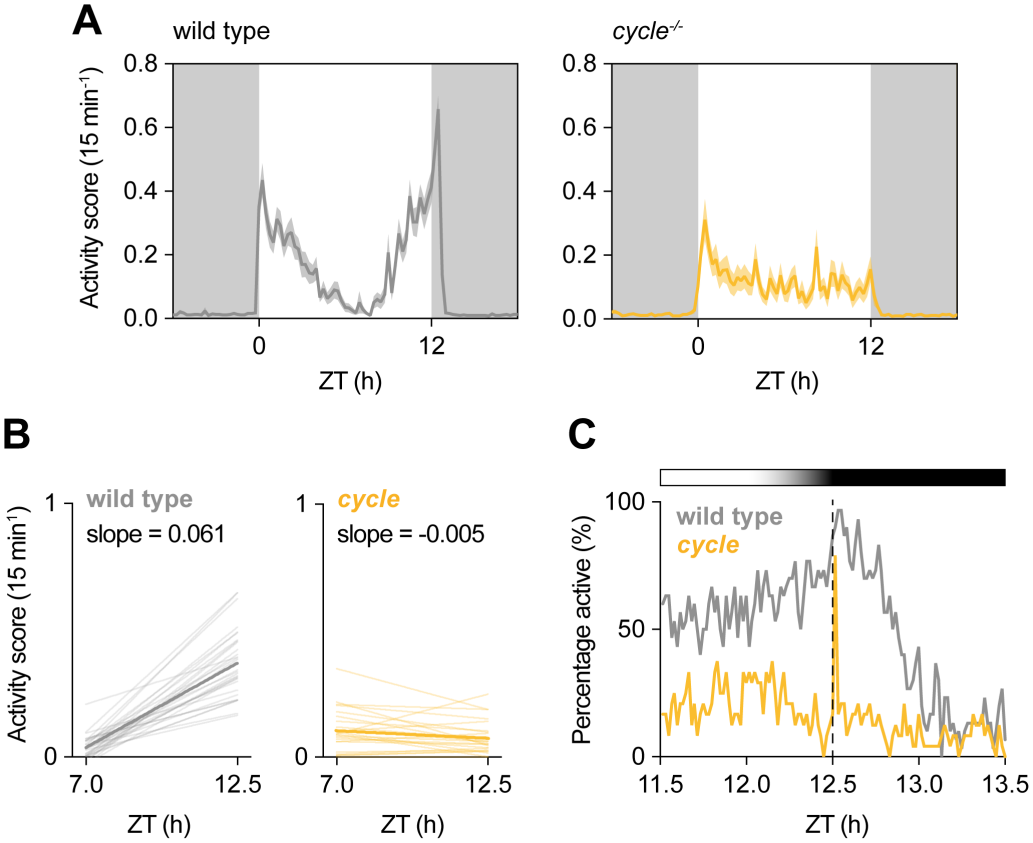
*Cycle* mutants show abolished temporal partitioning of daytime activity, related to Figure 4. (A) Locomotor activity rhythms of female mosquitoes under LD cycles. For all panels, data shown is the third day under LD cycles. For all panels, n = 30 and 24 animals for wild type and *cycle* mutants. White and grey shading represent light and dark phases. Line and shading show mean and SEM. (B) Linear regressions of activity score before light-off in wild type and *cycle* mutants. Thin lines show individuals and thick lines show group average. (C) Percentage of active animals around light-off in wild type and *cycle* mutants. Light and dark shading represents relative light intensity and dotted line represents lights-off.

**Figure S4.**
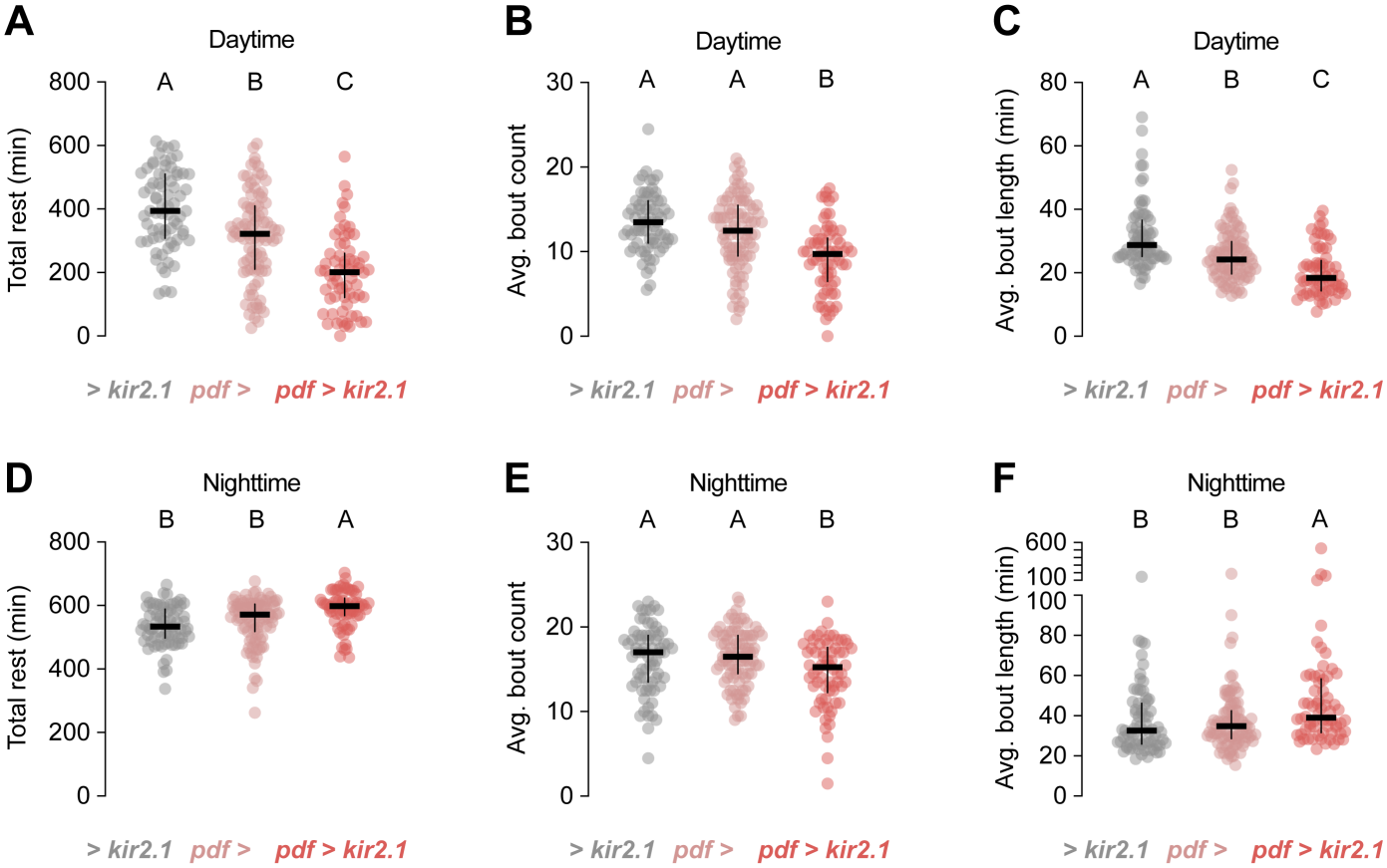
Rest metrics in *pdf >, > Kir2*.*1*, and *pdf > Kir2*.*1* genotypes, related to Figure 4. (A)Total daytime (ZT0–12) rest under LD cycles. For all panels, n = 71, 91, and 66 animals for *> Kir2*.*1, pdf >*, and *pdf > Kir2*.*1*, respectively. Rest is analyzed using activity recording of day 2–3 under LD cycles and defined as inactivity for > 10 mins. Kruskal–Wallis test with Dunn’s multiple comparisons. (B)Average daytime bout count under LD cycles. Kruskal–Wallis test with Dunn’s multiple comparisons. (C)Average daytime bout durations under LD cycles. Kruskal–Wallis test with Dunn’s multiple comparisons. (E)Total daytime (ZT0–12) rest under LD cycles. For all panels, rest is analyzed using activity recording of day 2–3 under LD cycles. Rest is defined as inactivity for > 10 mins. Kruskal–Wallis test with Dunn’s multiple comparisons. (F)Average daytime bout count under LD cycles. Kruskal–Wallis test with Dunn’s multiple comparisons. (G)Average daytime bout durations under LD cycles. Kruskal–Wallis test with Dunn’s multiple comparisons. Lines and error bars show mean and SEM and dots show individual animals. Data labeled with different letters are significantly different from each other (P < 0.05 in pairwise comparisons). Exact P values are provided in **Table S1**.

**Figure S5.**
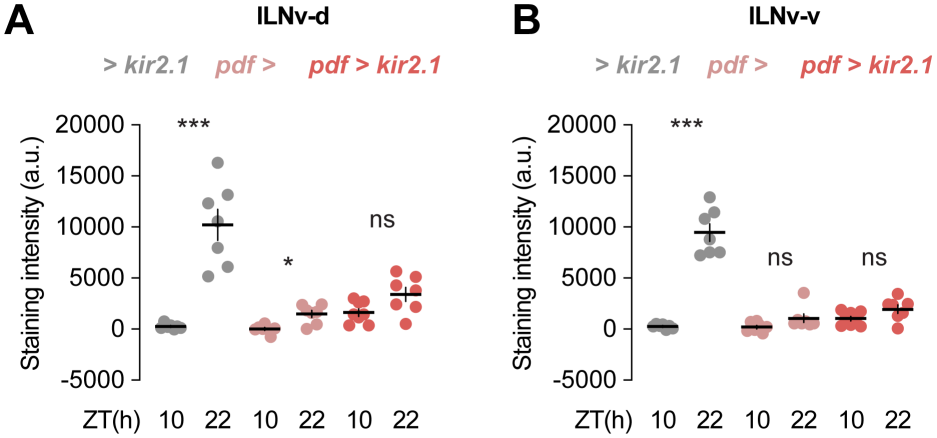
PER staining intensity in the lLNv-d (A) and lLNv-v (B) subgroups, related to Figure 5. Staining intensity reflects averaged measurements of somas within each subgroup (see Methods). For all panels, n = 6–8 brains for all genotypes and timepoints. Lines and error bars show mean and SEM and dots show individual animals. Mann-Whitney tests. *P < 0.05; ***P < 0.001; P > 0.05: not significant (ns). Exact n and P values are provided in **Table S1**.

